# A Bayesian model selection approach to mediation analysis

**DOI:** 10.1101/2021.07.19.452969

**Authors:** Wesley L Crouse, Gregory R Keele, Madeleine S Gastonguay, Gary A Churchill, William Valdar

## Abstract

Mediation analysis is a powerful tool for discovery of causal relationships. We describe a Bayesian model selection approach to mediation analysis that is implemented in our bmediatR software. Using simulations, we show that bmediatR performs as well or better than established methods including the Sobel test, while allowing greater flexibility in both model specification and in the types of inference that are possible. We applied bmediatR to genetic data from mice and human cell lines to demonstrate its ability to derive biologically meaningful findings. The Bayesian model selection framework is extensible to support a wide variety of mediation models.

## Introduction

Mediation analysis seeks to understand a causal process by determining whether an intermediate variable (M) explains (at least partially) the response of a dependent variable (Y) to changes in an independent variable (X). Though the causal interpretations of mediation analysis are subject to a number of assumptions [1], it has been widely used in both the social and natural sciences [2]. In a biomedical context, mediation analysis has been used to investigate how gene expression mediates the effects of genetic variants on complex phenotypes and disease [3, 4]. It has been used to infer causal relationships between biomolecular phenotypes, such as transcripts, chromatin states, and proteins [5, 6]. Here we focus on mediating genetic associations on biomolecular phenotypes, including gene expression, protein abundance, and chromatin accessibility, in samples of model organisms and human cell lines, but the mediation approach we introduce is broadly applicable.

Mediation analysis requires that X, M, and Y are measured in the same individuals, and its causal interpretation relies on assumptions that are not verifiable based solely on the data [7, 1, 8]. In particular, it assumes that the direction of causal effects is from X to Y (X*→*Y), referred to as the direct effect of X on Y, and from X to M to Y (X*→*M*→*Y), referred to as the indirect effect of X on Y. Also assumed is that these relationships are not subject to unobserved confounding, that observed confounders are conditioned on, and that observed confounders of M to Y do not depend on X. If these assumptions are satisfied, evidence for causal mediation lies in the magnitude of the indirect effect. If this indirect effect is non-zero, then M is a mediator of X on Y. Further, if M is a mediator and the direct effect is zero, then M is a complete mediator of X on Y, whereas if the direct effect is non-zero, then M is a partial mediator of X on Y. Our objective is to assess evidence of complete or partial mediation including the non-standard case where X comprises more than one independent variable.

Our motivating context is quantitative trait locus (QTL) mapping, where a univariate trait of interest Y (*e*.*g*., protein abundance) is associated with genetic variation represented in a matrix X, which could encode multiple variants or their combinations (*i*.*e*., haplotypes). In particular, we are interested in assessing whether one or more univariate candidate variables M may mediate the relationship between the genetic matrix and the trait. For example, a candidate mediator M could be the protein abundance of a gene encoded nearby a QTL for Y. In this case, it is reasonable to assume that the direction of causal effects is X*→*M*→*Y, and we further assume that the there is no unexplained confounding of these relationships. With the assumptions of mediation satisfied, our objectives are two-fold: 1) assess the evidence in favor of M being a causal mediator, 2) and determine if the mediation is partial or complete, given that X contains complex genetic information.

Traditional methods for mediation analysis are poorly suited to the dual objectives of detecting mediation and distinguishing complete from partial mediation when there are many candidate mediators and when X is a matrix. A classic approach for establishing mediation was introduced by Baron & Kenny [9]. This approach, termed the causal steps (CS) method, establishes evidence for partial or complete mediation by sequentially testing the relationships between X, M, and Y. Specifically, CS uses linear regression models to establish the following four conditions: 1) X has a marginal effect on Y [X*→*Y]; 2) X has an effect on M [X*→*M]; 3) M is a partial mediator of the effect of X on Y [M*→*Y|X]; and 4) M is a complete mediator of the effect of X on Y [X╨Y|M]. The CS method can accommodate a matrix of independent variables by using a likelihood ratio test for grouped predictors. Although the CS method is useful due to its conceptual accessibility, its implementation in a genomics setting with many candidate mediators can be awkward. In particular, it is not straightforward to combine statistics across the steps while also accounting for multiple testing, particularly for step (4), which requires a failure to reject a null hypothesis. This makes it difficult to succinctly summarize evidence for complete or partial mediation for many candidate M.

Other common tests for mediation analysis address the problems of the CS method by ignoring complete mediation and instead providing a single test statistic for the significance of the indirect effect. The indirect effect is formally given as the product of regression coefficients from X *→* M and M *→* Y | X, that is, the effect of M on Y controlling for the effect of X. Establishing that this coefficient product is non-zero gives evidence for (at least) partial mediation, but it does not provide information about complete mediation. The most popular methods for testing the indirect effect include the Sobel test [10], which is based on an approximation to the asymptotic distribution of the indirect effect; alternatively, bootstrapping can be used to assess significance [11], which does not make distributional assumptions but is computationally expensive. In addition to not providing information about complete mediation, the Sobel test does not generalize when X is a matrix rather than a vector.

Applications of mediation analysis to large scale genetic and molecular profiling data have used modified versions of the traditional tests described above. Approximations to CS have been used in the multi-parent Collaborative Cross (CC [12, 6, 13]) and Diversity Outbred mouse populations (DO [14, 5, 15, 16]). These studies identified QTL for gene expression (eQTL), protein abundance (pQTL), and chromatin accessibility (cQTL). Detection of a significant QTL for the target phenotype (Y) satisfies step (1) and detection of a QTL local to the molecular trait (M), *i*.*e*., near the genomic position of M, satisfies step (2). For a given phenotype QTL, a mediation scan is performed by testing the effect of X on Y as being mediated through each M (*e*.*g*., each observed gene transcript). For the approximation to CS, significant mediators are determined based on the reduction in log-odds (LOD) score before and after accounting for the effects of M (hereafter referred to as LOD drop). This approximates step (4) without requiring complete independence between X and Y. Notably, the LOD drop method does not directly check step (3) and, as a result, it may detect candidates that are correlated with the true mediator but are not mediating the effect of X on Y. Thus, care is needed when interpreting the LOD drop mediation scan.

More recent methodological developments in large-scale mediation of genetic and molecular profiling data include the multi-SNP intersection union test [17], an extension of the CS method that simultaneously models the overall effect of multiple genetic predictors by representing them as a similarity matrix-based (*i*.*e*., kernel-based) random effect, and the divide-aggregate composite null test [18], an extension of the joint significance test [19] that improves power relative to the Sobel and joint significance tests by utilizing an empirical null distribution. As with other methods based on the indirect effect, neither of these methods provide inference on distinguishing partial and complete mediation.

All of the approaches described rely on hypothesis testing, in which significance criteria are used to choose between nested alternative models. In our view, a more natural perspective for mediation analysis is provided by Bayesian model selection. Specifically, the goal of mediation analysis is to classify the relationship between X, M, and Y as a particular causal model. We note that the full space of potential causal models is not nested; the classification of the observed data into a particular causal model is, with finite data, necessarily uncertain; estimation of parameters when the model is uncertain ideally requires incorporation of model uncertainty into the estimate. These properties are well suited to the Bayesian model selection paradigm, which considers a set of potential models (nested or otherwise) and assigns to each a posterior probability.

Bayesian methods have been previously used in mediation analysis to estimate the posterior distribution of the indirect effect [20, 21, 22], and Bayesian model selection has been used to test for the presence of an indirect effect [23]. Here we develop a more expansive Bayesian model selection approach that considers the full space of causal models, distinguishing between—among others—complete mediation, partial mediation, and independence (where X affects M and Y independently, e.g. co-localized but distinct eQTL and pQTL). Bayesian approaches are often computationally intensive, and even infeasible for many applications. We avoid this problem by employing conjugate priors that allow us to avoid costly sampling when calculating posterior summaries. Our approach is computationally efficient, generalizes to multiple independent variables, and provides unique posterior summaries in the form of causal model probabilities.

## Results

We developed a Bayesian model selection approach, implemented in the bmediatR R package, to evaluate different causal models that define relationships between a continuous dependent variable Y, an independent (or exogenous) variable X, and a continuous mediator variable M. The causal models can be described as directed acyclic graphs (DAGs) (Figure 1). There are three possible edges (*a, b, c*) and we define each to be either present or absent using an indicator vector ***θ*** = (*θ*_*a*_, *θ*_*b*_, *θ*_*c*_), where for example ***θ*** = (1, 0, 0) denotes presence of *a* only. This leads to eight possible combinations of edges, each defining a different causal relationship (causal model) (ML1-8 in Figure 1). As an extension, the Bayesian model selection approach can also consider four additional causal models in which the direction of edge *b* (between M and Y) is reversed (ML9-12 in Figure 1, termed “reactive” models.), though some of these are are not causally identifiable (see Methods).

**Figure 1.**
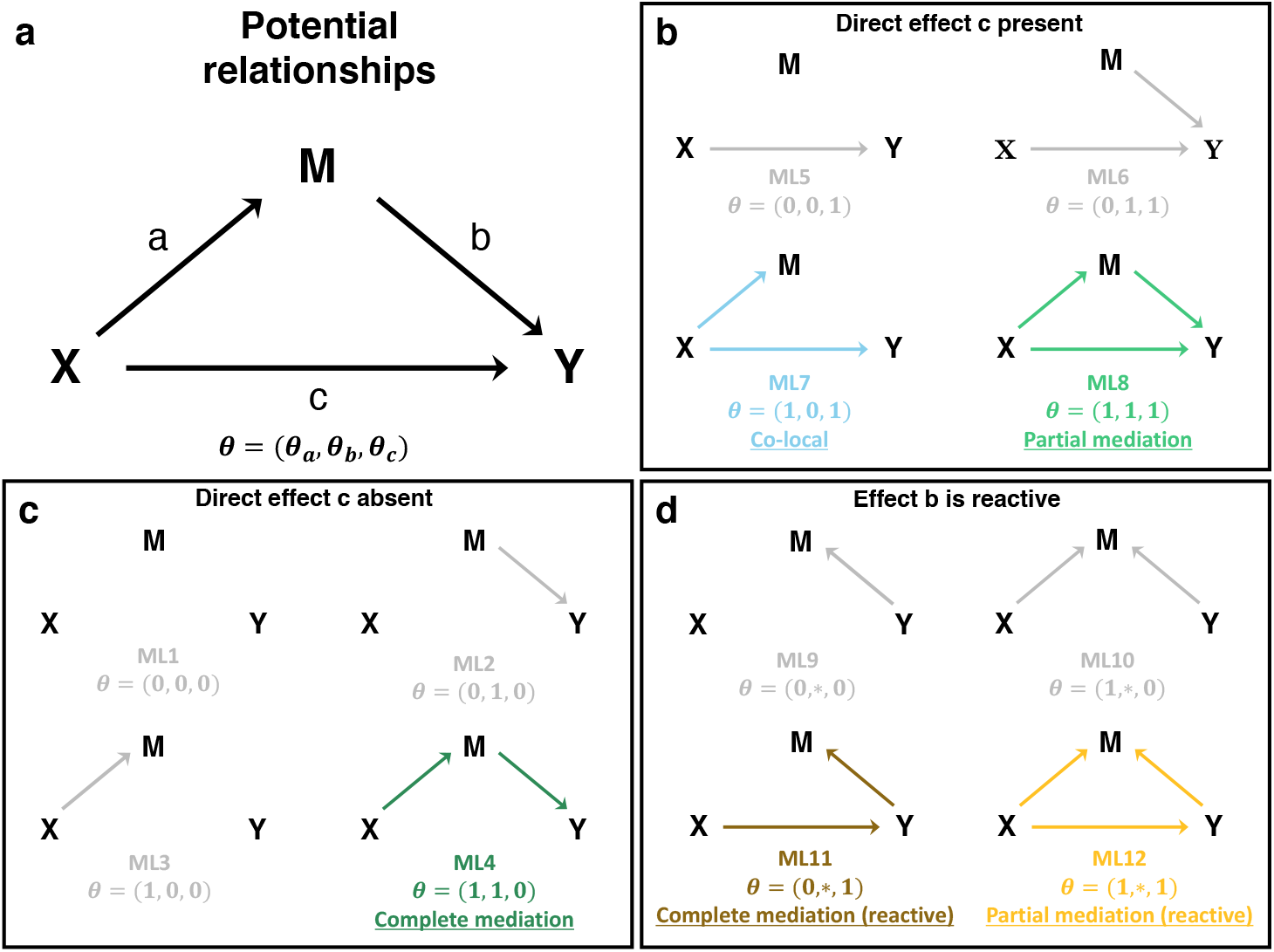
Possible relationships among X, M, and Y. X is assumed to be exogenous, and thus M and Y have no effects on X. A model and corresponding marginal likelihood (ML) are defined by the presence or absence of any the three edges *a, b*, and *c* according to an indicator variable ***θ***. In this work and by default in bmediatR, the direction of edge *b* is assumed to be from M to Y (M *→*Y), but a set of reactive models can also be accommodated in which the direction of edge *b* is reversed (M *←* Y), indicated with ***θ*** = (*θ*_*a*_, *, *θ*_*b*_). Models can be favored or even excluded by adjusting the model priors. By default, there are five models (ML1-3 and ML5-6) that represent non-mediation, *i*.*e*., the effect of X on Y, if present, is not mediated through M. The co-local model (ML7) represents a special case where there is no mediation between X and Y, but X independently affects M and Y. The complete mediation model (ML4) and the partial mediation model (ML8) represent cases where the effect of X on Y is explained, completely or partially, by the effect of X on M.

We are primarily interested in models that describe mediation, defined as any causal relationship where edges *a* and *b* are both present, *i*.*e*., ***θ*** = (1, 1, *θ*_*c*_), indicating a causal path from X to Y through M. Complete mediation describes models where X acts entirely through M and thus X conveys no additional information about Y beyond that provided by M (*θ*_*c*_ = 0); partial mediation describes models where X conveys additional information beyond that contained in M (*θ*_*c*_ = 1); and there is no mediation when *θ*_*a*_ = 0 or *θ*_*b*_ = 0, which includes the co-local model in which X affects both Y and M but independently, and also includes the model in which X and M affect Y but are independent of each other. Our approach calculates a posterior probability for each causal model ***θ*** given data X, M, and Y.

Bayesian model selection requires specifying prior probabilities for the models under comparison, as well as prior distributions for the parameters of those models (model priors). Priors can be used to incorporate external information into mediation analysis, which can improve power and reduce false positive rates. Since the results of Bayesian model selection analysis can be sensitive to the choice of priors, it is important to understand the role and interpretation of different prior specifications. The model priors are specified by assigning probabilities to each of the twelve possible configurations of ***θ*** (Figure 1). These probabilities can be viewed as weights and modified to define informative prior expectations on the relative frequency of different causal models. Setting a model prior to zero removes it from consideration, and this allows a user to define a set of allowable models. By default, we assign equal prior probability to the models that assume no reverse causality from Y to M for edge *b* (*i*.*e*., probability 1/8 for each model ML1-8 but probability zero for ML9-12).

Prior distributions on the effect sizes of the edges connecting X, M, and Y (effect priors) are given conjugate prior distributions, which yields a closed form for the joint likelihood of M and Y. The size of the effects is controlled by specifying the hyperparameters ***ϕ***, which are ratios of the edge effect sizes to the variability of the errors (Methods). The default specification ***ϕ*** = (1, 1, 1) assumes that all edges have *a priori* equal effect sizes, also equal in size to the error variances of M and Y. Using conjugate priors and specifying the edge effect sizes make computing the posterior fast and exact.

### Bayesian model selection performance in simulated data

We evaluated our Bayesian model selection against two alternative mediation analysis methods, the Sobel test and LOD drop, using simulated data. Data was simulated by first specifying X and then simulating M and Y according to linear models with normal error and effect sizes, expressed as proportion of variation explained, that ranged from 0.05 to 0.95. This approach allows X to represent a non-normal (and non-scalar) quantity. In particular, for genetics applications, X may represent minor allele counts, multi-state founder haplotype probabilities, or multiple genetic variants. Data were simulated for five scenarios: co-local, where X drives M and Y independently; partial mediation, where X drives Y both directly and indirectly through M; complete mediation, where X drives Y only through its effect on M; and reactive versions of partial and complete mediation, where X drives M through Y instead. For each scenario, we simulated 100 data sets with 200 observations each. For each simulated data set, we applied bmediatR’s default model priors (*i*.*e*., uniform over ML1-8 in Figure 1). We also considered two other types of model priors: reduced model priors, which sets a uniform prior probability over models that assume the direct effect *c* (ML4-8), and which is analogous to testing the indirect effect like the Sobel test; and expanded model priors, which sets a uniform prior over all models (ML1-12), including reactive ones (ML9-12) (Figures S1 and S2).

It should be noted that the Sobel test, LOD drop, and bmediatR provide different inferences. The Sobel test does not distinguish between partial and complete mediation, or their reactive forms. The LOD drop method does not distinguish between co-local, complete, partial mediation, or the reactive forms of the mediation models. Which models can be distinguished by the Bayesian model selection will depend on which prior model probabilities are non-zero, resulting in less or more complicated inference. If a direct effect from X to Y is assumed (edge *c* fixed), Bayesian model selection no longer distinguishes partial and complete mediation, similar to the Sobel test. Alternatively, more complex inference is possible by including reactive models as non-zero prior models in situations when the effect from Y to M is plausible, though inference between some models is not causally identifiable (see Discussion).

### Bayesian model selection performs well when genotypes are correctly specified

We evaluated the performance of the three mediation methods, with bmediatR set to default model priors, on data with a bi-allelic QTL with an allele frequency of 0.5, simulated under the co-local, partial mediation, and complete mediation models. Results from bmediatR with default model priors are shown in Figure 2; results from all prior settings and all simulation settings, including reactive, are shown in Figures S1 and S2.

**Figure 2.**
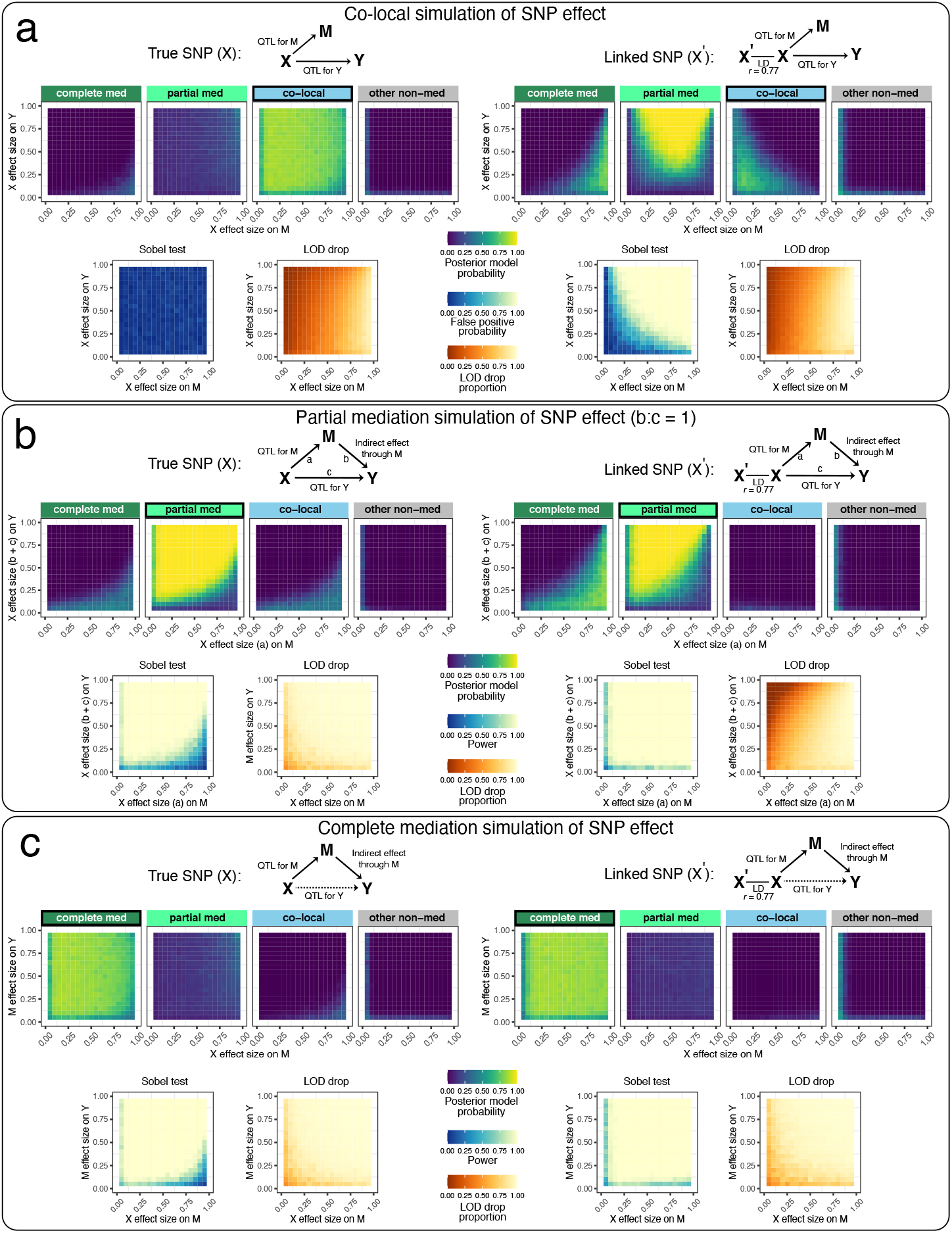
Performance of Bayesian model selection, Sobel test, and LOD drop in simulated data with a binary exogenous variable. Data for 200 individuals were simulated according to (a) co-local, (b) partial mediation, and (c) complete mediation models based on a balanced bi-allelic variant X. We applied mediation analysis with X as the true variant (left) and as a variant in linkage disequilibrium (*r* = 0.77) with the true variant (right). DAGs indicate the model used to simulate the data. Heat maps for Bayesian model selection represent the mean posterior probability associated with each inferred model for a range of fixed settings of the model parameters as indicated on x- and y-axes. Heat maps for the Sobel test represent false positive probability for co-local simulations and power for mediation simulations. Heatmaps for LOD drop represent mean LOD drop, scaled to the proportion of the simulated QTL’s LOD score. See Figures S1 and S2 for results from non-default model priors and all simulated models, including reactive.

The left panel of Figure 2a shows results for when the SNP affects both M and Y but independently, as would be the case for an eQTL and a pQTL that are co-local but otherwise unrelated. The top four boxes show the posterior probability given by bmediatR for four types of models: complete mediation (ML4), partial mediation (ML8), co-local (ML7), and other non-mediation models (ML1-3 and ML5-6). Each box is a heatmap of the posterior probability under all possible simulated effect sizes of X*→*M (x-axis) and X*→*Y (y-axis). Together, the four boxes show that, for all but the most extreme combinations of effect sizes, bmediatR overwhelmingly favors the (correct) co-local model. Below these are heatmaps for the Sobel test and the LOD drop. The Sobel test aims to identify mediation, and so any detections for this co-local scenario will be false positives. Its heatmap shows that the false positive rate is low in all cases. The LOD drop heatmap shows the magnitude of the signal for mediation by that method. LOD drop is seen to correctly register a minimal mediation signal when the effect size of the QTL for Y is large and that for M is small; but it incorrectly detects mediation when the QTL for M is large (*>*80%), a situation that is common in eQTL data.

The left panels of Figure 2b,c show results for the same set up but where mediation is present, either partial or complete. For partial (Figure 2b), bmediatR correctly identifies the type of mediation in most cases except for when the effect size of the QTL for M much is disproportionately large, in which case it is misclassified as complete or co-local. The Sobel test correctly identifies the presence of mediation (type unspecified) with a similar error pattern as for bmediatR. The LOD drop performs a little better than other methods, albeit with different error characteristics. For complete mediation (Figure 2c), the Sobel test and LOD drop perform slightly better than for partial mediation, while bmediatR accurately identifies the mediation type across all but the most extreme effect size settings.

### Misspecification of bi-allelic genotypes induces false mediation

Mediation was then evaluated when the bi-allelic X is misspecified. This can occur, for example, when X represents a variant that is imperfectly correlated with the true causal variant through linkage disequilibrium (LD) (Figure 2 right panels). In this setting, Bayesian model selection begins to favor the partial mediation model and the Sobel test begins to detect mediation when it is not present. LOD drop also suffers with misspecification, exacerbating its issues when the QTL on M is large. When data were simulated with mediation present (partial and complete), misspecification of X was less problematic (Figure 2b-c right), although Bayesian model selection was less accurate at distinguishing partial and complete mediation.

### Misspecification of multi-allelic genotypes as bi-allelic induces false mediation

Next we evaluated how mediation would be affected by misspecifying a multi-state haplotype effect. Data were simulated for 200 individuals with equal frequency of four alleles with distinct effects. Fitting X based on a bi-allelic SNP that tags the low and high haplotype groups worked well for complete mediation, but resulted in false mediation signals for co-local data, only correctly preferring the co-local model when at least one of the QTL on M and Y were small (*<* 50%) (Figure 3a). This issue was exacerbated when the SNP was more imbalanced, resulting in false mediation signal for co-local data across a wider range of effect sizes (Figure 3b). We then looked at simulated bi-allelic data but modeled the genetic effect as eight haplotypes. Bayesian model selection performed well for co-local and complete mediation data, aside from some edge cases with extreme effect sizes, most notably at the corners of large QTL on M with small QTL on Y (prefers complete mediation) and small QTL on M and large QTL on Y (prefers partial mediation) (Figure 3c).

**Figure 3.**
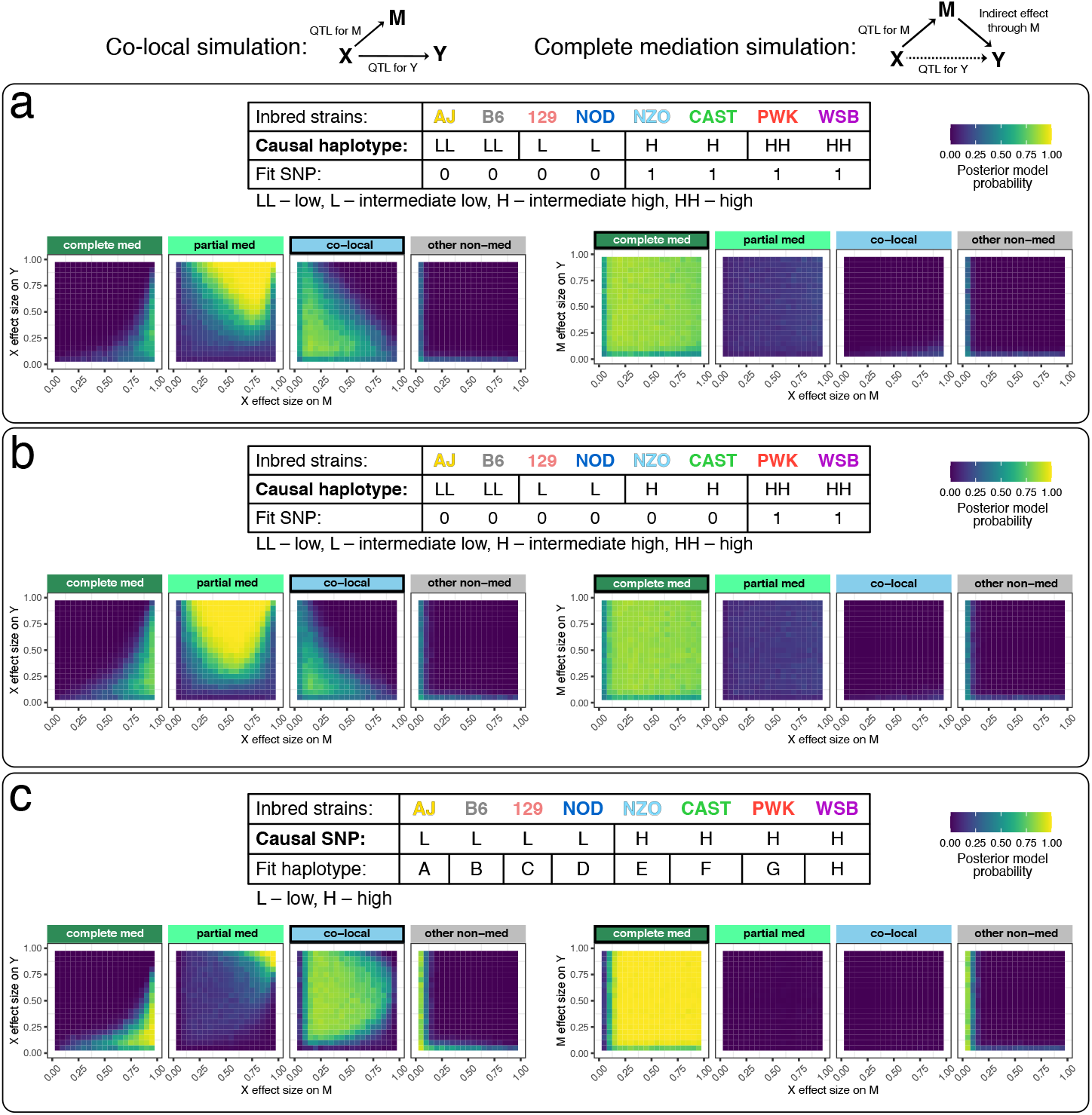
Performance of Bayesian model selection in simulated data with a multi-state exogenous variable. Data for 200 individuals were simulated according to co-local (left) and complete mediation (right) models. DAGs indicate the model used to simulate the data. (a-b) The genetic effect assumes four functional alleles with balanced allele frequencies (25%). Mediation analysis was performed using (a) a variant that tags the two higher functional alleles and (b) a variant that tags only the highest functional allele. (c) Data from a bi-allelic variant with allele frequency 50% were simulated, and mediation analysis performed using 8 founder haplotype states. Tables describe the relationships of the structure of X used to simulate the data versus the structure of X used in the mediation analysis. Heat maps for Bayesian model selection represent the mean posterior probability associated with each inferred model for a range of fixed settings of the model parameters as indicated on x- and y-axes.

### Bayesian model selection can detect mediation of multi-allelic QTL

To demonstrate mediation analysis in the context of multi-allelic QTL analysis, we simulated data based on the genomes of 192 DO mice [5]. A genetic locus X was randomly selected and M and Y were simulated assuming no mediation (QTL for Y, no QTL for M, M and Y uncorrelated; ML5), co-local (ML7), and complete mediation (ML4). QTL mapping was then performed for both M and Y to obtain LOD scores and estimated haplotype effects (Figure 4). The simulations demonstrate characteristic features of co-mapping QTL with correlated effects for co-local and complete mediation simulations, as they would appear in a multi-allelic QTL analysis. Data were simulated assuming a bi-allelic genetic variant, but the QTL mapping and Bayesian model selection were performed based on 8-state founder haplotypes.

**Figure 4.**
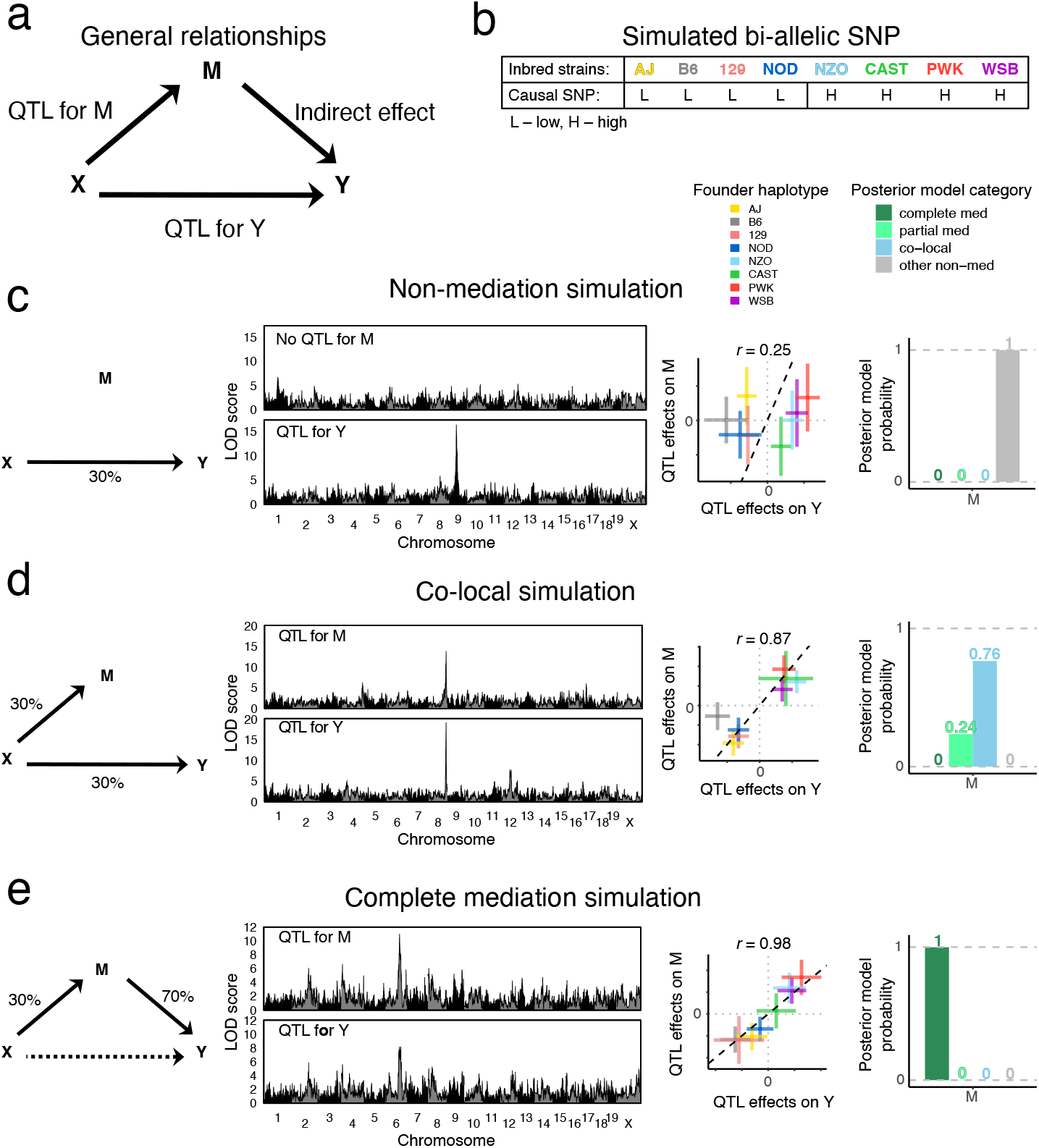
Illustration of Bayesian model selection applied to QTL mapping with simulated DO mouse data. (a) The DAG is labeled to indicate how each arm in the mediation model is interpreted in the QTL mapping setting. Y and M were simulated based on a bi-allelic QTL X at a randomly selected locus, with (b) each allele distributed to four founder strains. Genome-wide genotype data were obtained from 192 Diversity Outbred mice, according to one of three models: (c) M is a non-mediator of X on Y, (d) M and Y are independently driven by X (co-local), and (e) M is a complete mediator of X on Y, as illustrated with the corresponding model DAG (left). Genome-wide LOD scores for QTL mapping of M and Y, a scatter plot of the founder haplotype effects at the QTL for M and Y, and the Bayesian model selection posterior model probabilities are shown (from left to right).

### Bayesian model selection applied to DO mice

In order to illustrate how bmediatR works in QTL mapping applications, we analyzed previously reported liver proteomics data from 192 DO mice [5, 13]. We first illustrate how multi-state haplotypes help to identify the causal driver of a distal pQTL for *Snx4*. Then we look at a case in which bmediatR identifies a biologically plausible mediator for a distal pQTL of *Tubg1* where LOD drop favored a less plausible candidate.

### Multi-state haplotypes improves mediation inference for SNX4 distal pQTL

The *Snx4* gene is located on chromosome 16 and has a distal pQTL on chromosome 3 that co-maps with a local pQTL for *Snx7* (Figure 5). The proteins for both genes are sorting nexins that bind phospholipids, form protein-protein interactions, and play a role in membrane trafficking and protein sorting [24]. The haplotype effects of the *Snx7* local pQTL and the *Snx4* distal pQTL are highly correlated (*r* = 0.99) and reveal a complex haplotype effects pattern that cannot be explained by a single bi-allelic variant. We used the TIMBR software [25] to determine that the pQTL has as many as *≈*5 distinct functional alleles (Figure S3a). Bi-allelic variants in the pQTL region with alleles shared by the B6 and 129 strains partially match the haplotype effects and thus have strong associations with both proteins. In order to evaluate SNX7 as a mediator of the pQTL effect, we first applied Bayesian mediation using the peak SNP (with B6 + 129 allele) and find that this analysis assigns most of the posterior probability to the partial mediation model. In contrast, Bayesian mediation using the 8-state haplotype information finds strong support for complete mediation. We then evaluated all proteins (genome-wide) as potential mediators of the *Snx4* distal pQTL, and only SNX7 is identified as a likely complete mediator based on the log posterior odds (Figure 5e). The co-local model is highly unlikely for SNX7. Summing the posterior probabilities for complete and partial mediation we see clearly that SNX7 is the most likely candidate mediator of the distal QTL for *Snx4*.

**Figure 5.**
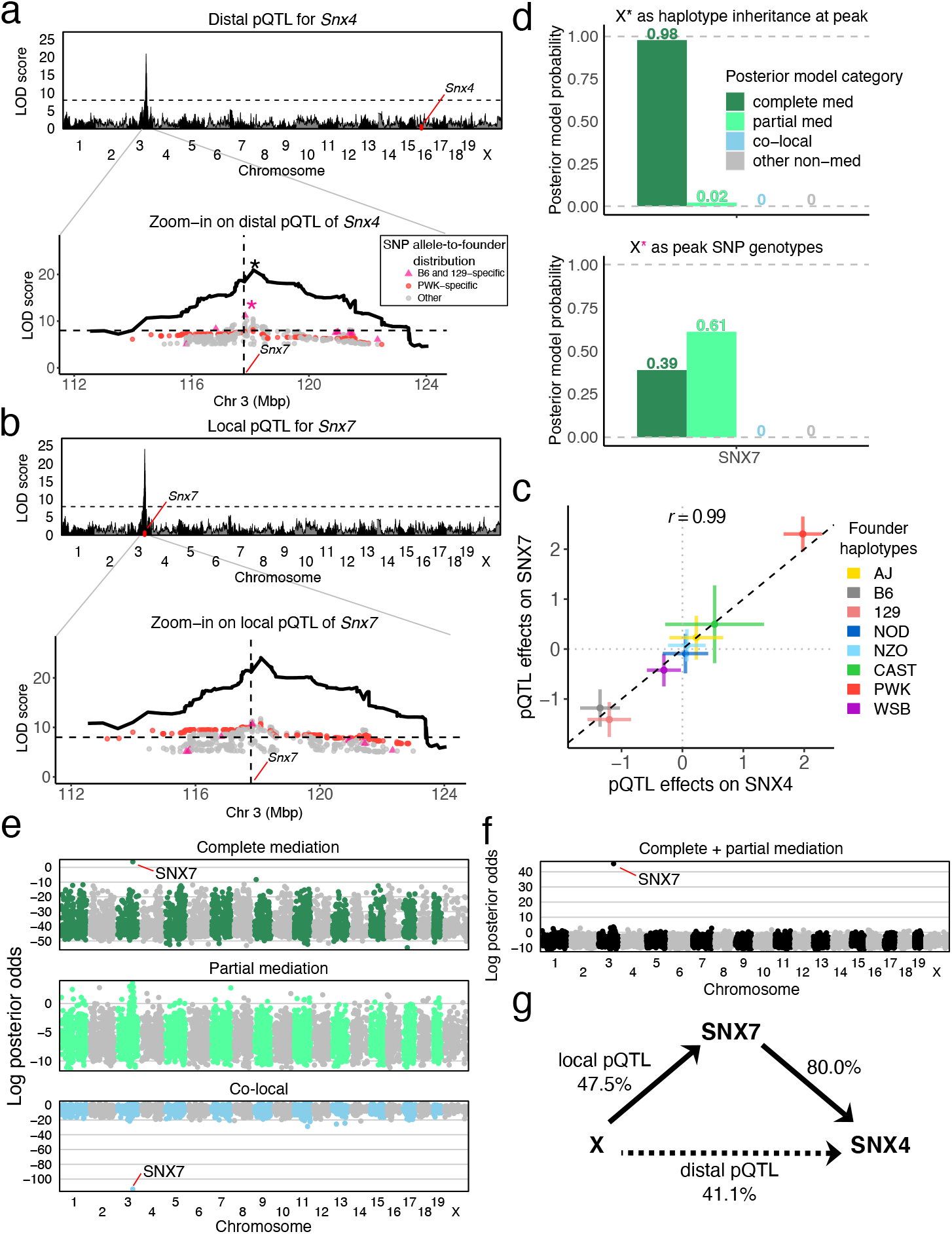
Mediation analysis of a distal pQTL for Snx4 in DO mice. Genome-wide LOD scores for associations of (a) SNX4 and (b) SNX7 abundance were performed using founder haploptye linkage mapping. Zooming into the QTL region, LOD scores for variant association within the pQTL region (peak *±* 5 Mbp) for bi-allelic vartiants with LOD scores *>* 5 are overlaid on the haplotype association LOD curve. Variants with alleles specific to B6 and 129 (pink) and PWK (red) are highlighted. (c) The founder haplotype effects at the pQTL are multi-allelic and highly similar for the two proteins. (d) Posterior probabilities of mediation models for the pQTL (top) using founder haplotypes and (bottom) using the peak bi-allelic variant. (e) Genome-wide mediation scan where all observed proteins are individually evaluated as mediators of the *Snx4* distal pQTL highlights SNX7 as a complete mediator and strongly indicates that the co-local model is unlikely. Each point represents the log posterior odds for a candidate mediator for the specified mediation model. (f) Summing complete and partial mediation posterior probabilties distinguishes SNX7 as the single best candidate mediator. (g) The complete mediation model with SNX7 as mediator of the *Snx4* distal pQTL is shown as a DAG with estimated effect sizes in units of percent variance explained. The dashed line indicates the strength of the distal pQTL that is not included in the model because it is completely mediated.

### Bayesian model selection favors a biologically plausible driver for TUBG1

The gene *Tubg1* is located on chromosome 11 and has a distal pQTL on chromosome 8. In our previous work, genome-wide mediation analysis by the LOD drop method identified NAXD as the best candidate mediator, but also revealed a significant LOD drop score for TUBGCP3 (Figure 6). Both mediation candidates have local pQTL. The *Naxd* local pQTL is stronger (LOD score *≈*40) than both the *Tubg1* distal and *Tubgcp3* local pQTL (LOD scores *≈*10). The pQTL haplotype effects for *Tubgcp3* and *Naxd* are highly correlated with the distal pQTL effects on TUBG1 (although flipped for NAXD) (Figure 6c). The allelic series could be either bi- or tri-allelic (*k* of 2-3; Figure S3b). We applied Bayesian model selection to both candidate mediators using the haplotype effects. The posterior probabilities at the two candidate mediators show that (Figure 6f) TUBGCP3 has 98% probability as a partial mediator, with the remaining probability for complete mediator. In contrast, NAXD, which is the best candidate using the LOD drop method, has 79% probability of being a complete mediator, 20% partial mediator, and 1% co-local probability. Due to the non-zero co-local probability of NAXD, TUBGCP3 is preferred based on the combined posterior odds of partial and complete mediation. A related tubulin gene, *Tubgcp2*, encoded on chromosome 7, has a distal pQTL at this locus on chromosome 8 (Figure 6g). Bayesian model selection analysis of the *Tubgcp2* distal pQTL also supports TUBGCP3 as a candidate mediator. These findings support a model for stoichiometric co-regulation of the protein constituents of the tubulin small complex [26, 27] that is mediated by the abundance of TUBGCP3.

**Figure 6.**
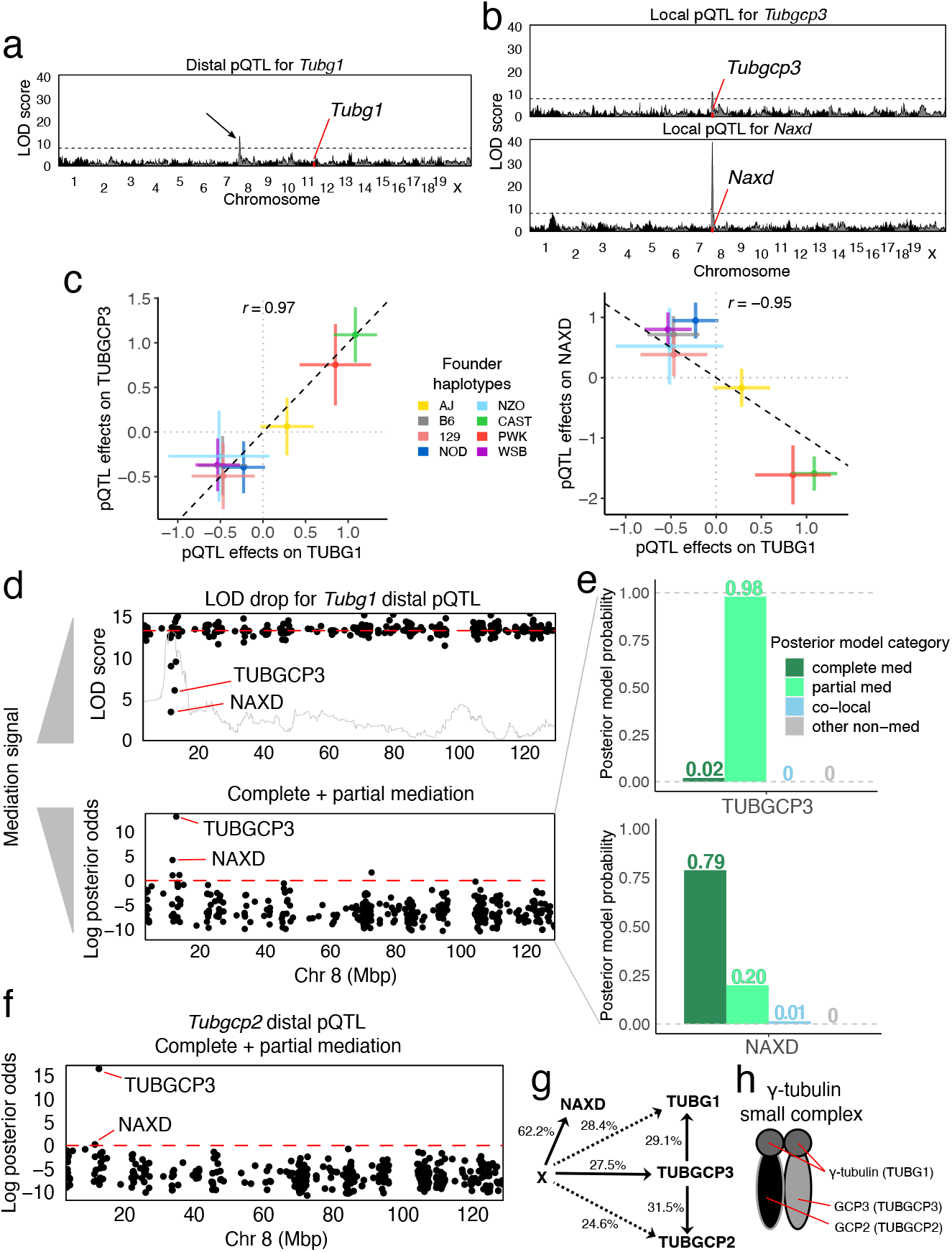
Mediation analysis of a distal pQTL for Tubg1 in DO mice. (a) Genome-wide LOD scores for TUBG1 abundance. Black arrow indicates distal pQTL on chromosome 8. (b) Genome-wide LOD scores for two genes *Tubgcp3* (top) and *Naxd* (bottom), with co-mapping local pQTL. (c) Comparison of the founder haplotype effects of the *Tubg1* pQTL with *Tubgcp3* (left) and *Naxd* (right) pQTL. (d) Mediation scans of all observed proteins on chromosome 8 by LOD drop with an overlay of the pQTL LOD scores in gray (top) and Bayesian model selection log posterior odds for mediation (bottom) show different prioritization for candidate mediators NAXD and TUBCP3. Note that low LOD drop scores indicate stronger mediation signal. (e) Posterior model probabilities for the *Tubg1* distal pQTL for candidate mediators (top) TUBGCP3 and (bottom) NAXD. (f) Mediation scan for the *Tubgcp2* distal pQTL identifies TUBGCP3 as the best candidate mediator. (g) The DAG summarizes the mediation analysis results with effect size estimates shown as percent variance explained. Dashed lines indicate the strength of distal pQTL effects that are not part of the model assuming complete mediation through TUBGCP3. (h) TUBG1, TUBGCP2, and TUBGCP3 comprise the *γ*-tubulin small complex.

### Mediation of genetic effects on gene expression through chromatin accessibility in human cell lines

We applied Bayesian model selection to data from 63 human lymphoblastoid cell lines (LCLs) using genotype [28], RNA-seq [29], and DNase-seq [30, 31] data. We used both Bayesian model selection and the Sobel test to identify chromatin regions near transcription start sites that may act as mediators of gene expression. Previously, genetic variants that affect chromatin accessibility and gene expression were mapped in the LCLs [30]. Variants associated with chromatin accessibility were also likely to be associated with the expression of nearby genes, indicating that chromatin accessibility may be a common mechanism by which transcription is regulated.

To illustrate, we looked at the expression of gene SLFN5 which was strongly associated with local genetic variation (eQTL). These genetic variants were also strongly associated with chromatin accessibility (cQTL) (Figure 7a). The SNP most strongly associated with SLFN5 expression and chromatin accessibility is located within an interferon-stimulated response element (ISRE) in the first intron of SLFN5 [30], and the co-mapping chromatin site is directly above the SNP. The juxtaposition of chromatin site to gene makes it a likely mediator for the expression of SLFN5, an interferon-regulated gene, because it controls the accessibility of the ISRE to transcription factors [32]. Bayesian model selection and the Sobel test both detect partial mediation.

**Figure 7.**
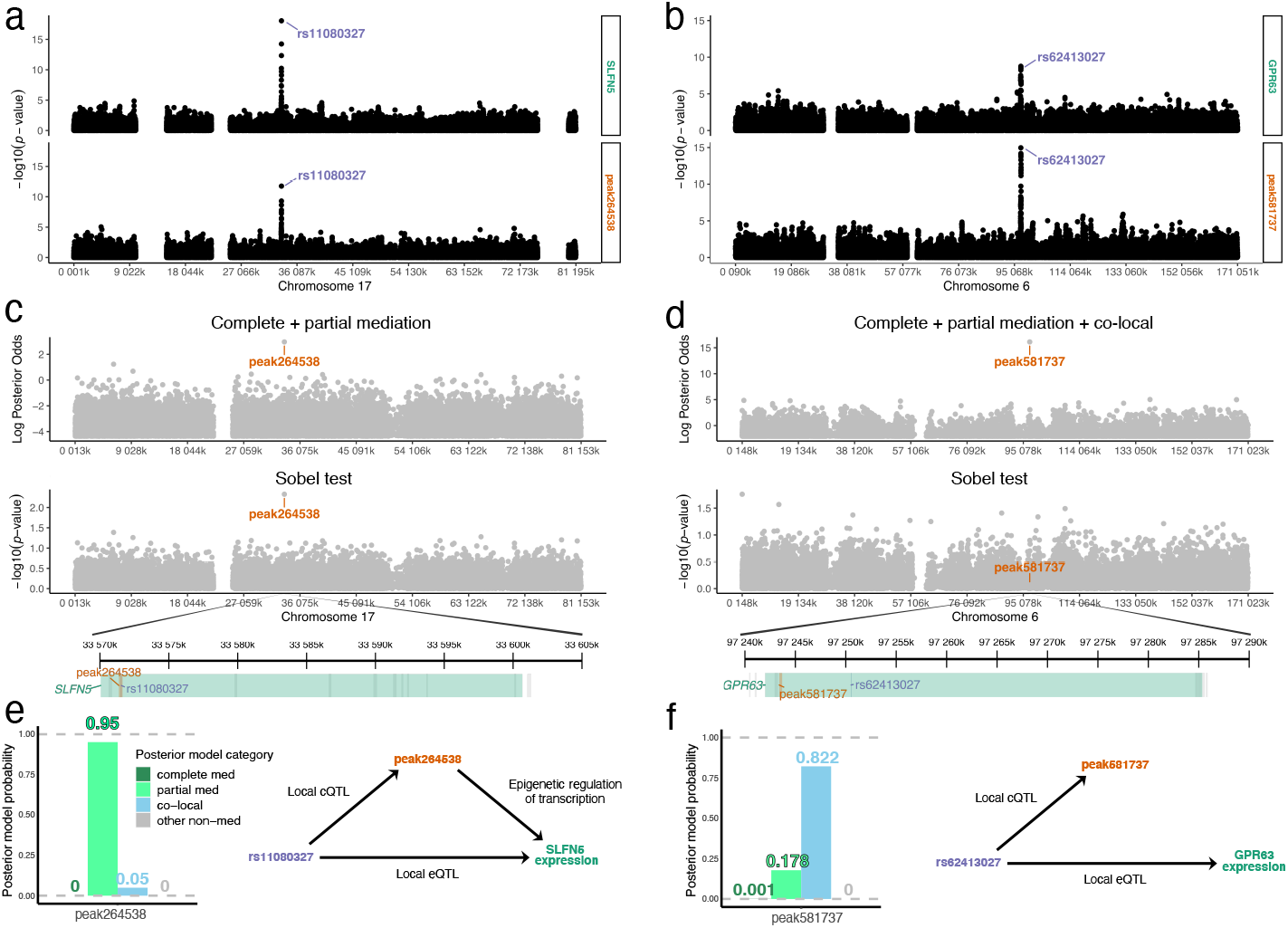
Mediation analysis of local chromatin state and gene expression data in human cell lines. SNP associations with (a) SLFN5 and (b) GPR63 expression (top) and nearby chromatin accessibility (bottom) for variants on the genes’ chromosome. Peak SNPs are labeled. Mediation results for the (c) SLFN5 eQTL and (d) GPR63 eQTL. Log posterior odds from Bayesian model selection (top), −log10 *p*-values from Sobel test (middle), and zoomed-in window highlighting gene start, peak SNP, and peak mediator (bottom). Peak mediator or co-local chromatin peak is labeled. Each gray point represents a chromatin peak candidate mediator located near the gene of interest. For SLFN5 expression, complete and partial mediation models were summed in the posterior summary from Bayesian model selection. For GPR63 expression, the co-local model was also summed with the mediation models. Posterior model probabilities from Bayesian model selection for the peak mediator and co-local chromatin peaks and the implied DAG for the (e) SLFN5 eQTL and (f) GPR63 eQTL.

Co-mapping eQTL and cQTL can also represent independent signals. A SNP within 10 Kbp of the start of GPR63 is a local eQTL and a local cQTL for a nearby chromatin site (Figure 7b). The Sobel test does not support the chromatin site as a mediator of the genetic effect on GPR63 expression. Bayesian model selection results are consistent with the Sobel test in ruling out mediation but, unlike the Sobel test, it clearly identifies the relationship as co-local.

## Discussion

Results from our Bayesian model selection analysis can be sensitive to the choice of model prior; therefore, it is important to understand the role and interpretation of different prior specifications. In our framework, is is possible to define any configuration and relative weight of causal models using the model prior. For example, if one believes that X is not a QTL for most candidate mediators, we could assign a smaller prior probability to models that include the *a* edge; or, if we believe that reactive models are unlikely but do not want to exclude them completely, we could assign these models a small prior probability (with the caveat that some reactive models are not identifiable). The appropriateness of such choices depends on the context of the analysis and the set of candidate mediators under consideration (*e*.*g*., considering only genes nearby X, versus all genes in the genome).

A straightforward elaboration of our approach is to use a set of candidate mediators to learn an empirical prior distribution over the causal models. If suitably constructed, an empirical prior could improve power within a given dataset. An empirical prior could be implemented, for example, by computing the model posterior for all candidate mediators using the uniform model prior, and then taking the average over all posteriors as the empirical prior distribution for downstream analyses. Conceptually, this approach is similar to the empirical null employed in Liu *et al* [18]. As an illustration, in the case where X is a QTL for Y, if most of the candidates are not actually mediators, an empirical prior would put a high prior probability on models that include the *c* edge and do not represent mediation. Our simulations indicate that the default uniform model prior led to reasonable model selection inference over a range of true effect sizes and causal models. That said, exploring alternative prior distributions over the causal models is an intriguing avenue for future research, and many strategies could be implemented within our flexible framework.

There may also be opportunities to improve inference by adjusting the effect size hyperparameters, ***ϕ***. By default, we assume *a priori* equal effect sizes for each edge, which are also equal to the size of the error variances. Tuning these hyperparameters to reflect prior beliefs about relative effect sizes may improve posterior model inference, provided these prior beliefs are closer to the truth. It is possible to put prior distributions on the effect size hyperparameters, but doing so would complicate posterior inference, as conditioning on these hyperparameters makes inference fast and exact. Other possibilities for setting the effect size hyperparameters include maximum *a posteriori* estimation (*e*.*g*., via grid search) or defining an empirical prior based on the data, though we have not explored these ideas here.

Mediation analysis critically assumes that there is no confounding between M and Y, *i*.*e*., M *←* U *→* Y. In the presence of confounding, M and Y may be associated even if there is no direct effect from M to Y. Known confounders can be adjusted for as covariates (**Z** in the Methods). Unobserved confounders present a more fundamental problem. Approaches that account for unobserved confounders could be implemented, but only as part of a two-stage approach, where latent factors are inferred first, naive to the mediation model, and then either removed or included as covariates in the mediation analysis. In our real data applications, we analyzed experimental data from mice and human cell lines, which we assumed was unconfounded after controlling for sex, batch, and other experimental variables.

There are other non-mediation approaches that are commonly used to investigate the relationship between X, M, and Y. These include Mendelian randomization (MR) [34, 35, 36, 37], which is an instrumental variables approach for estimating the causal effect of M on Y in the presence of confounding. MR involves building a predicted M (using X *→* M), and regressing Y on the predicted (rather than observed) M. This approach is robust to confounding, but it requires the additional strong assumption of no direct effect of X on Y (*i*.*e*., no partial mediation). This means that classical MR cannot be used to distinguish between partial and complete mediation. Emerging MR methods relax the assumption of no direct effect and may be useful in this context, but this field is rapidly developing and a full review of these methods is beyond the scope of this paper. A different, non-causal method for evaluating the relationship between M and Y is colocalization analysis [38, 39, 40, 41, 42], which evaluates if two variables (M and Y) share a common X, when only one of many independent variables are under consideration. This approach is typically motivated by fine-mapping genetic variants (*i*.*e*., evaluating candidate X, rather than candidate M), and it does not establish a causal relationship between M and Y in the way that mediation analysis can.

Mediation analysis commonly assumes that the direction of causal effect is from M to Y. This assumption must be justified based on the context of the analysis. For example, the mediator of a distal pQTL is expected to possess a corresponding local QTL. In the presence of reverse causality from Y to M, however, standard mediation analysis (*i*.*e*., inference that excludes the reactive models) will still detect an edge from M to Y, yielding spurious evidence for mediation. In our framework, is it straightforward to include the reactive models in settings where reverse causality is a possibility. We examined inference in the presence of reverse causality via simulation (Figures S1 and S2). These results demonstrate that it is possible to distinguish complete mediation from complete reactive mediation, but also indicate that it is not possible to distinguish partial mediation from reactive partial mediation. This is not unexpected, as these models are not identifiable. Our simulation results also show that, for the priors we specified, true partial mediation can appear more consistent with reactive complete mediation than partial mediation, for a particular space of effect sizes (Figure S2b bottom row). These findings emphasize the importance of the assumption that M *→* Y in mediation analysis. By default, our bmediatR software excludes reactive models, and we recommend interpreting inferences made using the reactive models with caution.

Bayesian model selection provides an opportunity for more flexible model specifications than are possible for other methods, such as the Sobel test. In particular, the exogenous variable X can be categorical (with more than two groups) or multivariable. This is useful for modeling genetic effects in some contexts, as we demonstrated by the improvement in mediation using 8-state haplotypes in the DO mouse data. Mediation analysis with multiple exogenous variables [43] is easily implemented in bmediatR. The bmediatR software offers a framework that could be generalized to account for moderated mediation [44, 9, 45], in which the effect of the mediator is moderated by another factor. Increasing the complexity of the model space can increase computation time and may require more complex posterior summaries. Despite these hurdles, compared to methods like the Sobel test, implementing more general mediation models settings is straightforward within the Bayesian model selection framework.

Lastly, we caution that mediation analysis is sensitive to misspecification. In our simulations, we showed that when a genotype X is misspecified (*i*.*e*., a haplotype is incompletely described by a single variant), a true co-local model can be inferred as mediation. Simulations in other studies have demonstrated that greater precision in the measurement of Y relative to M can lead to true complete mediation appearing as partial mediation [46]. These issues highlight the need for careful examination and independent validation of inferences based solely on mediation analysis.

## Conclusions

Mediation analysis is a powerful tool for discovery and integrative analyses of high dimensional biological data, with the caveat that proper caution and awareness of the potential pitfalls are needed. Here we describe a flexible Bayesian model selection approach to mediation analysis that is implemented as the R package bmediatR. Using simulations, we show that bmediatR performs as well or better than established methods including the Sobel test, while allowing greater flexibility in both model specification and in the types of inference that are possible. We applied bmediatR to genetic data from mice and from human cell lines, demonstrating its ability to derive biologically meaningful findings. The Bayesian model selection approach provides a flexible framework to support further advances in mediation analysis methods.

## Methods

### The standard mediation model, the Sobel test, and LOD drop

The standard mediation model can be described using two linked linear models. For individual *i* = 1, …, *N*, let *y*_*i*_ be the value of the primary outcome or dependent variable, let *m*_*i*_ be the value of the mediator variable, and let *x*_*i*_ be a (scalar) independent variable. Mediation analysis frames the relationships between these variables in terms of the following regressions:

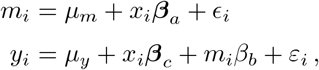

where *µ*_*m*_ and *µ*_*y*_ are intercepts, *∊*_*i*_ and *ε*_*i*_ are normally distributed, independent random noise variables, *β*_*a*_ is the effect of *x*_*i*_ on *m*_*i*_, *β*_*b*_ is the effect of *m*_*i*_ on *y*_*i*_, and *β*_*c*_ is the effect of *x*_*i*_ on *y*_*i*_. The effects *{β*_*a*_, *β*_*b*_, *β*_*c*_*}* correspond to edges *{a, b, c}* in the DAG in Figure 1. In the language of mediation, *β*_*c*_ describes the direct effect of *x*_*i*_ on *y*_*i*_, whereas the combination of *β*_*a*_ and *β*_*b*_ describe the indirect effect of *x*_*i*_ on *y*_*i*_ via *m*_*i*_.

The Sobel test for mediation focuses on the estimation of the product *β*_*a*_*β*_*b*_. This product, which provides a single number description of the indirect effect, can take any value from *−∞* to *∞* but is zero only when either *β*_*a*_ = 0 or *β*_*b*_ = 0 or both. The traditional implementation of the Sobel test uses classical estimates of *β*_*a*_*β*_*b*_, and bootstrapping or other approximations [47], to estimate confidence interval for *β*_*a*_*β*_*b*_ and thereby a *p*-value for *β*_*a*_*β*_*b*_ ≠ 0. Bayesian implementations of the Sobel test analogously determine a posterior for *β*_*a*_*β*_*b*_ and thereby a suitable tail probability [20] or Bayes factor [23]. In either implementation, the Sobel test provides a succinct way to quantify the evidence for mediation and uncertainty about the strength and direction of the indirect effect. By contrast, the CS approach [9], and thereby its approximation, the LOD drop method (both defined in the Introduction), is awkwardly constructed [48], lacks power [19, 49], and does not quantify uncertainty; it does, however, take into account the X *→* Y relationship, potentially enabling discrimination between partial and complete mediation. Note that distinguishing partial and complete mediation is sensitive to misspecification [46] and its value is subject to debate [50].

### Mediation with multiple independent variables

The standard mediation model is trivially adapted to cases where the independent variable is multivariable, that is, of dimension *D >* 1: the scalar predictor *x*_*i*_ and scalar effects *β*_*a*_ and *β*_*c*_ are simply replaced by their *D*-vector counterparts, **x**_*i*_, ***β***_*a*_, ***β***_*c*_. In this case, the CS (or LOD drop) approach to testing mediation may still be applied but, because the product ***β***_*a*_*β*_*b*_ is a vector, the Sobel test cannot. Sobel test-like procedures have been developed for specific cases, including a bootstrap-based test for when X is multicategorical [43] and a bespoke Bayesian model for when X is a scalar-by-multi-category interaction [21], but these are computationally intensive and do not generalize easily to multivariable X.

### A Bayesian model selection approach for mediation analysis

We propose a Bayesian model selection approach that combines the generality of the CS method with the inferential coherence of the Sobel test. Our approach is most similar to the one in Nuijten et al. 2015 [23], which is also a Bayesian formulation of mediation, but only considers a single independent variable in X. To describe our approach, we first elaborate the standard mediation model to

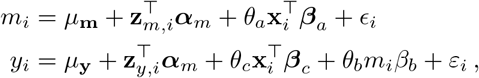

where **x**_*i*_ is a *D*-length column vector, ***β***_*a*_ and ***β***_*c*_ are *D*-length effects vectors, **z**_*m,i*_ and **z**_*y,i*_ are column vectors encoding any covariates for *m*_*i*_ and *y*_*i*_, ***α***_*m*_ and ***α***_**y**_ are the corresponding covariate effects, 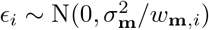 and 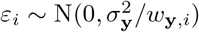 are noise variables with individual-specific weights *w*_**m**,*i*_ and *w*_**y**,*i*_, and ***θ*** = *{θ*_*a*_, *θ*_*b*_, *θ*_*c*_*} ∈ {*0, 1*}*^3^ are indicator variables denoting the presence or absence of edges {*a, b, c*} in the DAG in Figure 1, such that, for example, ***θ*** = {1, 1, 0} denotes complete mediation (*a* and *b* active) with no effect of edge *c*. The parameter space of ***θ*** contains 2^3^ = 8 possible combinations of edges, each of which corresponds to a particular causal model. From a Bayesian perspective, making inferences about the identity of the true causal model is equivalent to calulating the posterior distribution of ***θ***, that is,

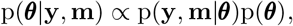

Where 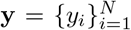 and 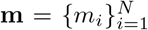. This involves calculating a joint likelihood for Y and M given a causal model, p(**y, m**|***θ***), and a prior distribution over causal models, p(***θ***). These are described separately below.

### Likelihood and priors conditional on a causal model

The conditional joint likelihood function is given by

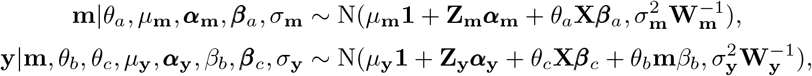

where **Z**_**m**_ and **Z**_**y**_ are matrices for the covariates, 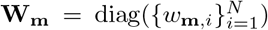 and 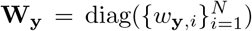 are diagonal matrices of observation weights. By default, **Z**_**m**_ = **Z**_**y**_ = **0**, and **W**_**m**_ = **W**_**y**_ = **I**.

Conjugate priors are given for the following variables:

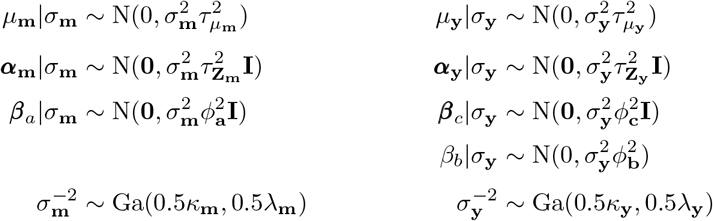

These variables can be integrated from the likelihood due to conjugacy, giving a closed form expression for the marginal joint likelihood function:

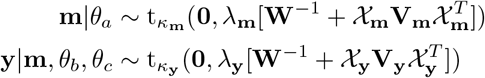

where *𝒳*_*m*_ and *𝒳*_**y**_ are concatenated design matrices and **V**_**m**_ and **V**_**y**_ are prior covariance matrices for the effect variables, specifically,

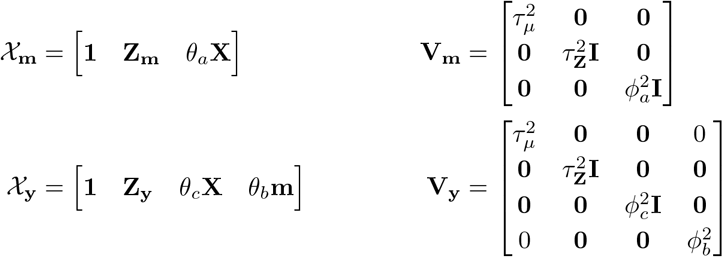

This marginal joint likelihood is evaluated for all causal models ***θ*** = (*θ*_*a*_, *θ*_*b*_, *θ*_*c*_), given prior hyperparameters ***κ*** = (*κ*_**m**_, *κ*_**y**_), 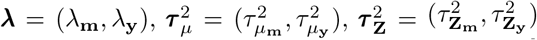, and 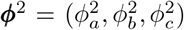. Non-informative priors are used for the scale of the data [***κ*** = ***λ*** = (0.001, 0.001)], the location of the intercept [***τ*** _***μ***_ = (1000, 1000)], and covariate effects [***τ*** _**Z**_ = (1000, 1000)]. The hyperparameter ***ϕ***^2^ controls the prior effect size of each edge, relative to error, on **m** or **y**. We set ***ϕ***^2^ = (1, 1, 1) by default, such that effect sizes for all edges are equal and relatively large *a priori*.

### Priors for the causal models

In our framework, a prior probability is assigned to each possible causal model. The default model prior places uniform prior probability across the eight models for which edge *b* is either absent or present as M*→*Y (no reactive models). That is,

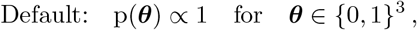

such that the prior probability for each of ML1-8 in Figure 1 is 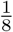. We also consider two other prior specifications: the reduced model prior, which additionally assumes that edge *c* is present, that is,

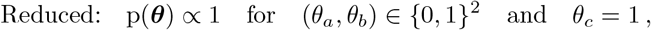

such that the prior probability for each of ML4-8 is 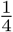; and the expanded model prior, which allows models with a reversed edge *b* (*i*.*e*., Y*→*M) and is described in more detail below. In our framework and using our bmediatR software, it is possible to specify priors as any distinct set of allowable causal models, or, with greater granularity, as relative weights across the set of models. This feature provides a foundation that could accommodate empirically-determined prior weights out-of-the-box.

### Allowing reverse causality between M and Y

The bmediatR software can also model reverse causality between the candidate mediator and trait (*i*.*e*., Y*→*M). This can be described using a third state for *θ*_*b*_, denoted by *θ*_*b*_ = *\**, giving a total of twelve possible causal models encoded by ***θ***. These include the eight causal models previously described, as well as four additional reverse causality (or “reactive”) cases, given by (*θ*_*a*_, *θ*_*b*_ = *\*, θ*_*c*_), where models with *θ*_*b*_ = *\** are specified with the roles of all Y- and M-related variable above interchanged (explicit formulae given in Supplemental Methods). Although any set of relative prior probabilities could be used, we here consider a simple prior (the expanded prior) that is uniform over the 12 causal models, that is,

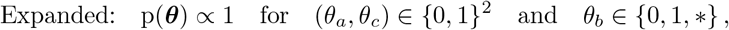

such that the prior probability for each of ML1-12 in Figure 1 is 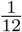.

Note that two pairs of these twelve causal models are not identifiable from one another: ***θ*** = (0, 1, 0) is indistinguishable from ***θ*** = (0, *\**, 0); and ***θ*** = (1, 1, 1) (partial mediation) is indistinguishable from ***θ*** = (1, *\**, 1) (reactive partial mediation). The marginal joint likelihood functions for these pairs of models are identical, and these causal relationships cannot be distinguished without additional information. Despite this limitation, we include all twelve causal relationships as possibilities in our expanded model prior, and emphasize that inference about the direction of causality in these cases depends entirely on prior information and assumptions.

### Simulation of QTL data

To simulate data with an exogenous variable representing genetic variation (*e*.*g*., SNPs, founder haplotypes), we expanded our previous approach for simulating QTL in the Collaborative Cross mouse population [51]. The model describes relationships between two traits with shared genetic drivers, representing either co-local or mediation.

We simulate a single trait with a single QTL based on a simple linear model:

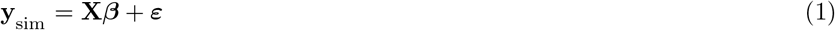

where **y** is the trait vector, **X** is the design matrix of the genetic information (*e*.*g*., SNP allele count, founder haplotype count), ***β*** is the genetic effects vector (*e*.*g*., SNP allele effect, haplotype effects), and ***ε*** is a random noise vector. We scale ***β*** and ***ε*** such that their relative contributions to **y**_sim_ match a specified proportion of variation of **y**_sim_ explained by **X*β***, *i*.*e*., the effect size of X on Y. If an initial genetic effect vector (***β***_raw_) is not specified, we sample one according to ***β***_raw_ ~ N(**0, I**), where **I** is the identity matrix of rank equal to length of ***β***. An initial noise vector is also sampled as ***ε***_raw_ ~ N(**0, I**_*n*_) where **I**_*n*_ is the identity matrix of rank *n*, the number of rows of **X**. For a specified effect size *φ*^2^ (ranging from 0 to 1), we define scaling factors for ***β***_raw_ and ***ε***_raw_ to approximate the desired effect size of X on Y:

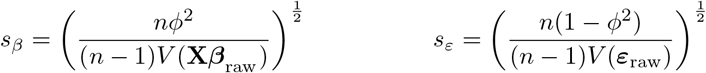

where *V* () returns the population variance of its argument. The effects and noise vectors are then scaled: ***β*** = *s*_*β*_***β***_raw_ and ***ε*** = *s*_*ε*_***ε***_raw_.

For the co-local simulations, both the dependent variable **y**_sim_ and the mediator variable **m**_sim_ are simulated as in Equation 1 with the same ***β***_raw_ but independent draws of ***ε***_raw_. For the complete mediation simulations, **m**_sim_ is simulated according to Equation 1, and the dependent variable **y**_sim_ is simulated based on a linear model with the mediator as a predictor:

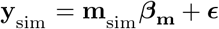

where ***β***_**m**_ and ***E*** are derived from scaled raw variables just as ***β*** and ***ε*** were in Equation 1 in order to strictly control how M contributes to variation in Y (*i*.*e*., effect size of M on Y).

The partial mediation simulations are necessarily more complicated. The mediator **m**_sim_ is again simulated according to Equation 1. The linear model for **y**_sim_ includes effects from both X and M:

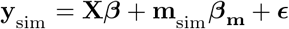

where ***β*** is the direct effect of X on Y and ***β***_**m**_ is the indirect effect. ***β, β***_**m**_, and ***E*** were derived from scaled raw variables as before in order to strictly control how X and M together contribute to variation in Y (*i*.*e*., combined effect size on Y). Reactive partial and reactive complete mediation were simulated exactly as partial and complete mediation, but with **y**_sim_ and **m**_sim_ swapped in the mediation procedures.

For the large-scale simulations of a single local (Figures 2, 3, S1, S2), **X** represented balanced functional allele counts (SNP or founder haplotypes) for 200 individuals. One hundred simulations were performed for each combination of effect size on M and Y (ranging from 0.05 to 0.95 at regular intervals of 0.05) for each data-generating model (co-local, partial mediation, complete mediation, reactive complete mediation, and reactive partial mediation). For the partial mediation simulations, the ratio of direct and indirect effect were fixed at 1 (b:c = 1). For each data-generating model, Bayesian model selection, Sobel test (if **X** represented a SNP), and LOD drop were summarized. For the simulations demonstrating multi-allelic QTL analysis with mediation in Figure 4, the design matrix **X** represents additive effects of founder haplotypes at a randomly sampled genetic locus from a population of 192 DO mice [5]. The underlying simulated genetic effect was from a bi-allelic SNP, with each allele present in four of the founder strains.

### Diversity Outbred mouse data

The DO mouse data represent 192 animals [5] with both gene expression and protein abundance from bulk liver tissue, representing a subset of a larger cohort of 850 animals [52]. Approximately equal numbers of males and females are present as well as animals on standard and high fat diets. The data are publicly available for interactive analysis as a QTL Viewer (https://github.com/churchill-lab/qtlapi) which also allows a bulk download of the underlying R data files (https://qtlviewer.jax.org/viewer/SvensonHFD).

### QTL analysis

Samples of liver tissue were collected and processed for quantitative massspectrometry as previously described [5]. Estimation and normalization of the protein abundance data from component quantitative peptide data and subsequent QTL analysis have been previously described [13]. Genetic mapping was based on final quantities output from the rank-based inverse normal transformation (RINT) [53].

The examples related to the distal pQTL of *Snx4* and *Tubg1* were previously identified [13]. QTL were mapped using the following linear mixed effect model:

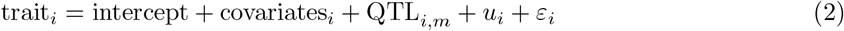

where trait_*i*_ is the phenotype of interest (abuandance of a protein) for individual *i*, intercept is a shared intercept that represents the mean trait value, covariates_*i*_ is the effect of known covariates on individual *i*, QTL_*i,m*_ is the effect of genetic variation at genomic interval *m* on individual *i, u*_*i*_ is a random error term that accounts for the similarity of individual *i* to other samples proportional to overall genetic relatedness (kinship effect), and *ε*_*i*_ is the unstructured error on individual *i*. For the QTL term: 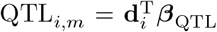, where **d**_*i*_ is the vector of additive dosages of founder haplotypes for the genomic interval *m* and ***β***_QTL_ are the haplotype effects at the putative QTL, estimated as fixed effects. The random error terms are modeled as **u** ~ N(**0, K***τ* ^2^) and *ε*_*i*_ ~ N(0, *σ*^2^), where **K** is the realized genetic relationship matrix excluding information from the chromosome of the current genomic interval *m* (“loco” method), and *τ* ^2^ and *σ*^2^ are variance components for *u*_*i*_ and *ε*_*i*_, respectively. Covariates adjusted for include sex, diet, and DO litter (2 levels). Equation 2 is fit at genomic intervals spanning the entire chromosome, comprising a QTL genome scan. Models were fit using the qtl2 R package [54]. For the variant association performed for the *Snx4* and *Snx7* pQTL, Equation 2 was used, but with the QTL_*i,m*_ adjusted. Instead of representing the effects of doses of founder haplotypes, variant allele dosages were imputed based on the founder haplotype dosages and the variant genotype-to-founder strains distribution (SQLite variant database: https://doi.org/10.6084/m9.figshare.5280229.v3).

### Haplotype effect estimation

To compare the similarity of the genetic effects on M and Y, the correlation coefficients between haplotype effects of QTL were calculated. To stabilize haplotype effect estimates, instead of the fixed effect estimate from Equation 2, they were estimated as best linear unbiased predictions (BLUPs) in the qtl2 R package.

### Modeling the allelic series of QTL with TIMBR

We modeled the allelic series at the pQTL for to SNX4 and TUBG1 using a Bayesian hierarchical model as implemented in TIMBR [25]. The model is roughly equivalent to Equation 2, with the kinship effect excluded in order to make the computation feasible. A Chinese restaurant process prior was used for the allelic series, as well as the prescribed shape and rate parameters of 1 and *≈*2.33, respectively, for the concentration parameter, which favors smaller numbers of functional alleles with low variance.

TIMBR models founder haplotype uncertainty at the QTL based on the 36 genotype states. We reconstructed founder haplotype probabilities from the available genotype array data (https://www.jax.org/research-and-faculty/genetic-diversity-initiative/tools-data/diversity-outbred-database), which represented 187 of the 192 mice.

### Human cell line data

The genotype data from 119 Yoruba LCLs represent variants from the intersection of HapMap2 and HapMap3, coded as the major allele count for each SNP [28]. The RNA-seq data [29] were adjusted with WASP [55] and normalized [30], representing 69 LCLs [28]. Both genotype and RNA-seq data are publicly available for download (http://eqtl.uchicago.edu/jointLCL/). The DNase-seq data for the 69 cell lines with RNA-seq data were used as previously processed [31]. The overlap of samples across genotypes, RNA-seq, and DNase-seq was 63 LCLs.

### Chromatin accessibility and expression QTL analysis

To identify candidates for co-mapping eQTL and cQTL, we correlated the expression of gene with chromatin peaks within 80 Kbp of the gene start. QTL mapping was then performed for genes and chromatin peaks that had correlations *>* 0.5. Mapping was performed by regressing a trait (gene expression or chromatin accessibility) onto the genotypes of individual SNPs located on the gene’s chromosome, and compared to a null model with no genotype term. QTL were called for SNPs that produced a −log10 *p*-value *>* 8, representing a stringent threshold for 63 individuals. Gene-chromatin peak pairs were then filtered to those with an eQTL and cQTL co-mapping to the same SNP. For each passing gene, all chromatin peaks on the gene’s chromosome were tested as candidate mediators using both Bayesian model selection, without default prior settings, and the Sobel test.

## Availability of data and materials

All data and R code used to generate the results are available at figshare (https://doi.org/10.6084/m9.figshare.14912484).

## Acknowledgments

We thank Michael I. Love of the University of North Carolina at Chapel Hill for discussions related to this work and for pointing us to publicly available human cell line data.

## Authors’ contributions

WLC, GRK, and GAC conceived the project. GAC and WV supervised and directed the research. WLC developed and implemented the methodology. WLC and GRK developed the software package. GRK and MSG analyzed data and visualized results for the manuscript. WLC, GRK, and MSG drafted the manuscript. GAC and WV edited the manuscript. All authors read and approved the manuscript before review.

## Supplemental Methods

### Joint likelihood for causal models in the reactive mode

For causal models describing reverse causality between M and Y, that is, the reactive models ***θ*** = (*θ*_*a*_, *θ*_*b*_ = *\*, θ*_*c*_), the roles of M and Y are switched relative to the other models, so that the conditional joint likelihood p(**y, m**|***θ***) in these case is

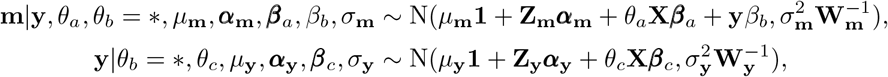

where *β*_*b*_ is now the scalar effect of **y** on **m**. Prior distributions for all variables are unchanged (except for ***θ***). When *θ*_*b*_ = *\**, the marginal joint likelihood function is given by:

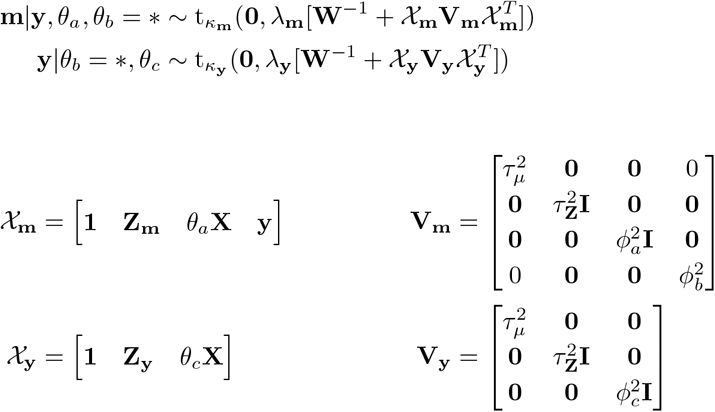

Hyperparameters for *κ, λ*, ***τ*** _*µ*_, ***τ*** _**Z**_, and ***ϕ***^2^ are unchanged.

## Supplemental Figures

**Figure S1.**
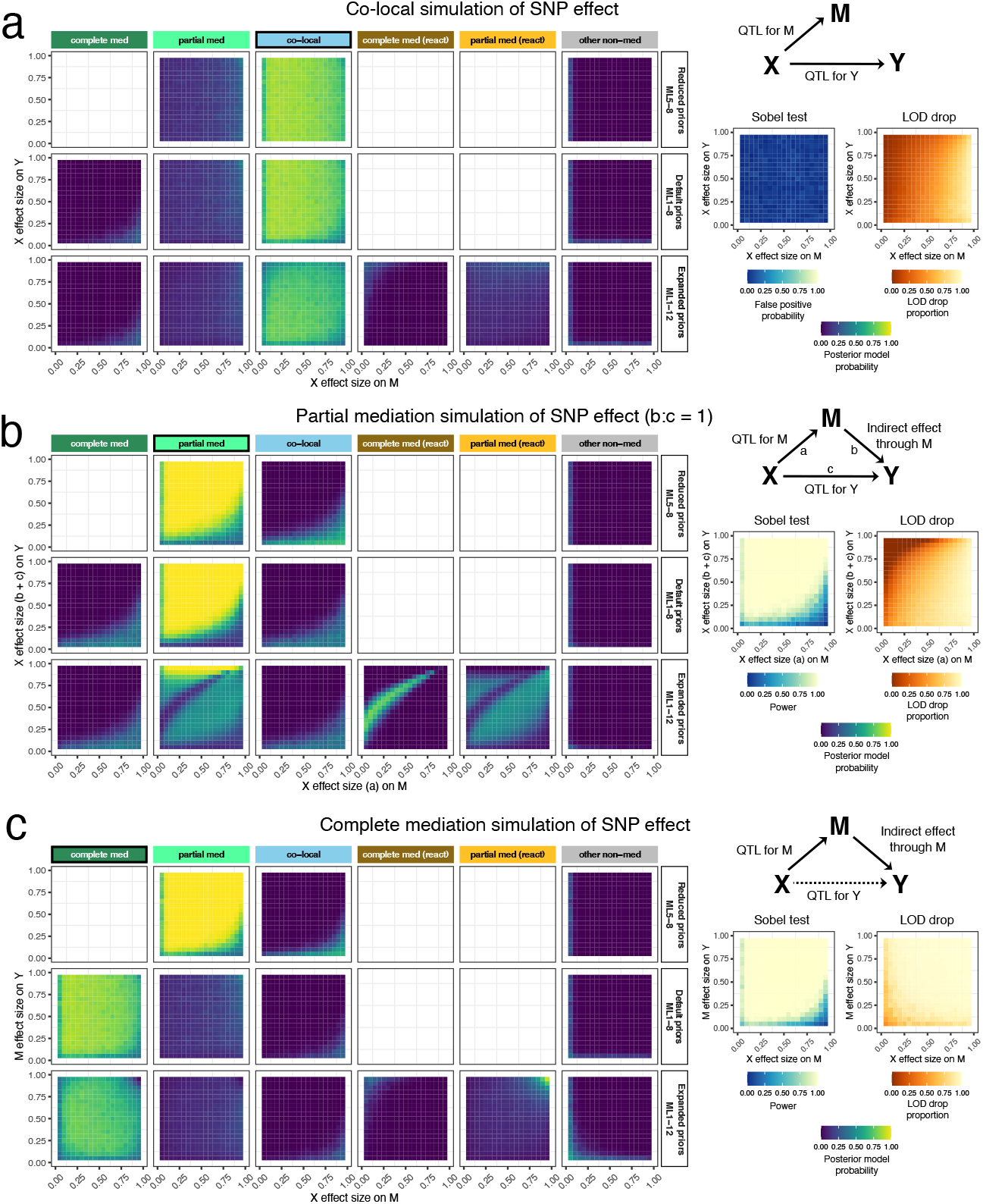
Performance of Bayesian model selection, Sobel test, and LOD drop in simulated QTL data, including reactive models. Data for 200 individuals were simulated according to (a) co-local, (b) partial mediation, and (c) complete mediation models from a balanced bi-allelic SNP. There is no misspecification of X. The DAGs represent the underlying models of each set of simulations. Heat maps for Bayesian model selection represent the mean posterior probability associated with each inferred model for a range of fixed settings of the model parameters as indicated on x- and y-axes. Heat maps for the Sobel test represent false positive probability for co-local simulations and power for mediation simulations. Heat maps for LOD drop represent mean LOD drop, scaled to the proportion of the simulated QTL’s LOD score. Empty squares represent posterior model categories not evaluated based on the set of admissible models encoded in the model priors.

**Figure S2.**
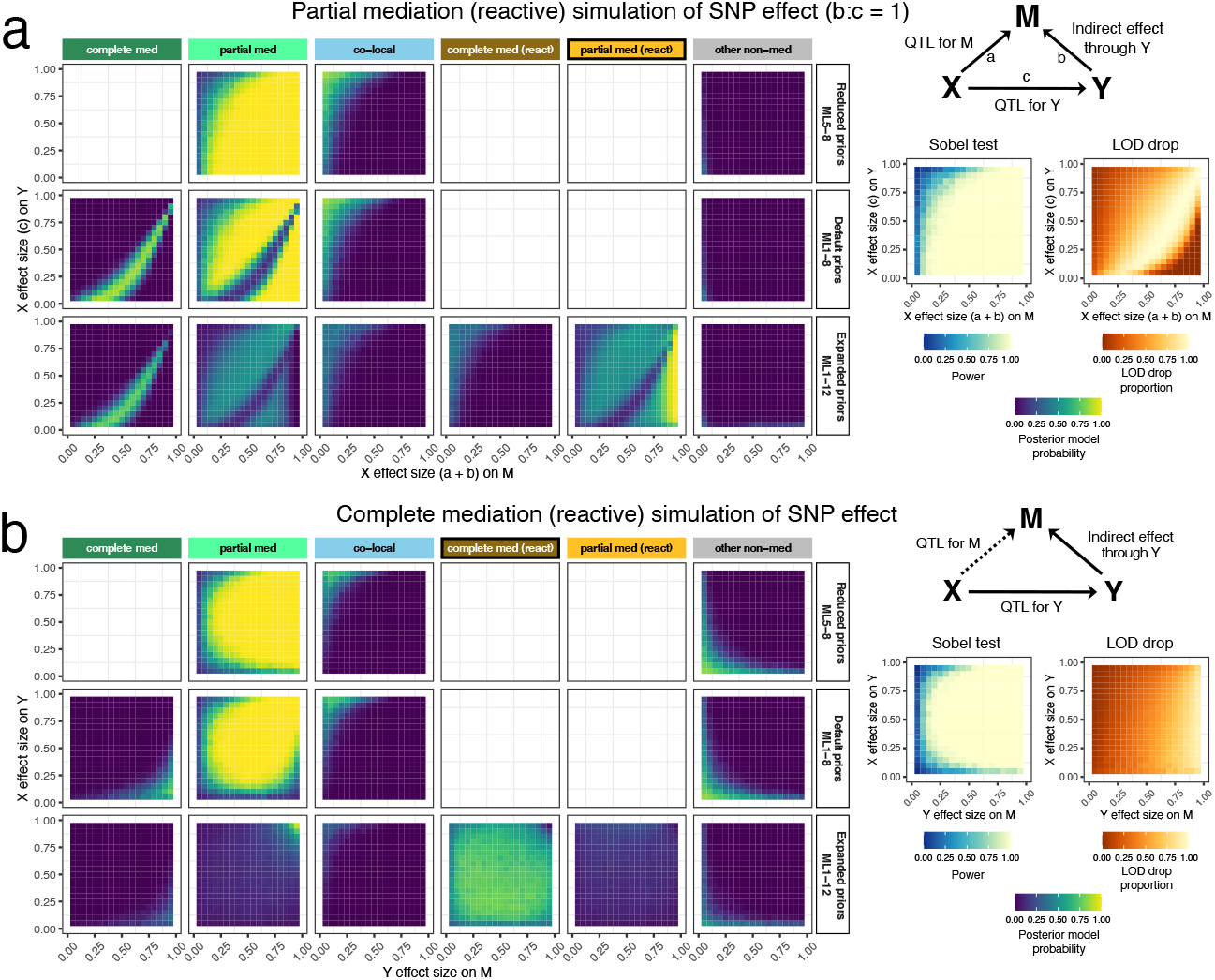
Performance of Bayesian model selection, Sobel test, and LOD drop in simulated reactive QTL data, including reactive models. Data for 200 individuals were simulated according to (a) reactive partial mediation and (b) reactive complete mediation models from a balanced bi-allelic SNP. There is no misspecification of X. The DAGs represent the underlying models of each set of simulations. Heat maps for Bayesian model selection represent the mean posterior probability associated with each inferred model for a range of fixed settings of the model parameters as indicated on x- and y-axes. Heat maps for the Sobel test represent false positive probability for co-local simulations and power for mediation simulations. Heat maps for LOD drop represent mean LOD drop, scaled to the proportion of the simulated QTL’s LOD score. Empty squares represent posterior model categories not evaluated based on the set of admissible models encoded in the model priors.

**Figure S3.**
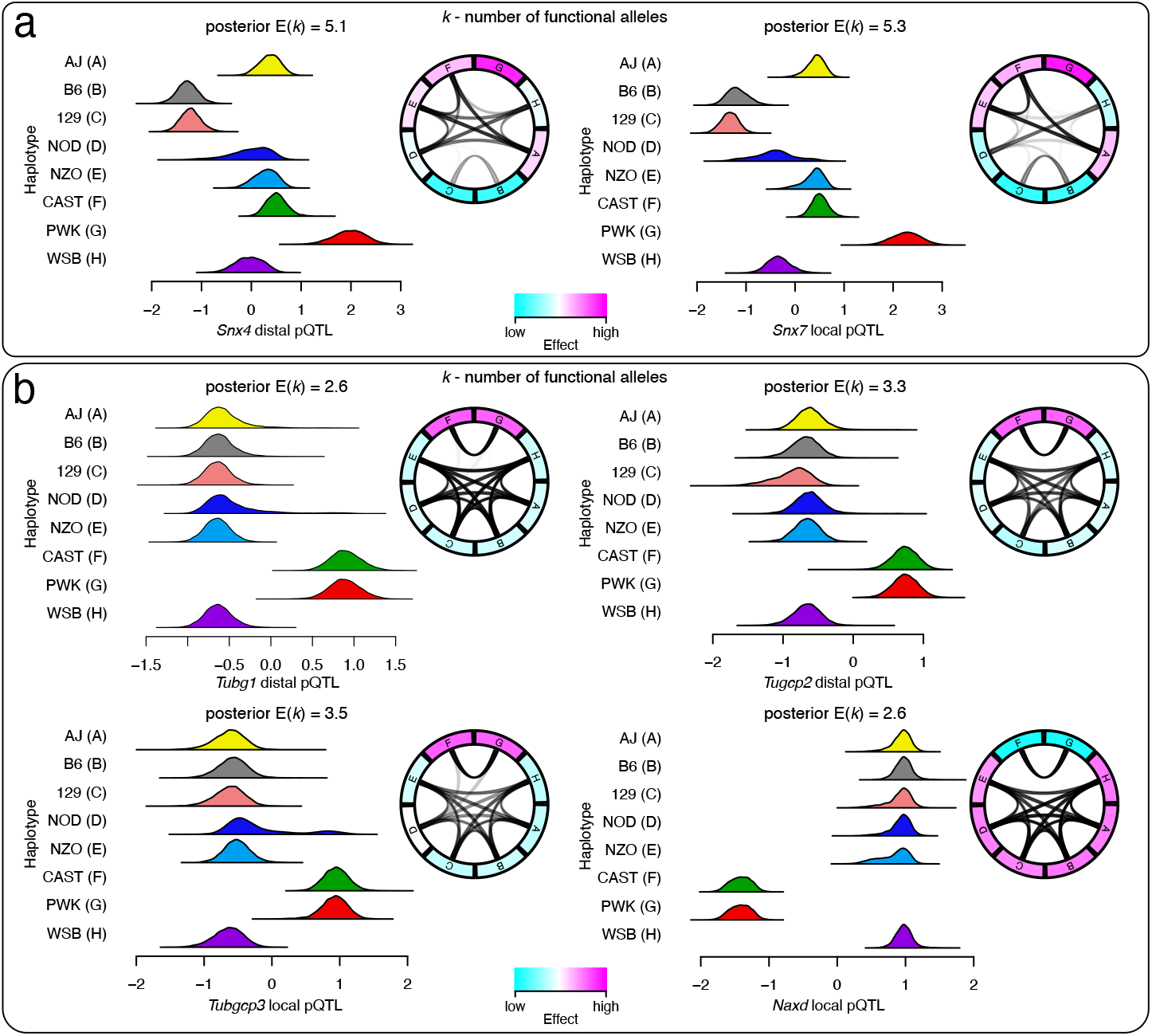
Modeling the allelic series of pQTL for Snx4, Tubg1, and related genes identified through mediation. The allelic series were modeled with TIMBR. (a) Posterior haplotype effects and allelic series for the *Snx4* distal pQTL and the local pQTL for its mediator, *Snx7*. Haploytpe effects are represented as histograms of the posterior samples. Allelic series are represented as circos plots where the opacity of the links represent how often TIMBR assigned founder haplotypes to the same functional allele. (b) Posterior haplotype effects and allelic series for the *Tubg1* and *Tubgcp2* distal pQTL and the local pQTL for their candidate mediators, *Tubgcp3* and *Naxd*. The posterior expected number of functional alleles, *k*, is included for each pQTL.

